# Gene identity, not recoding strategy, determines the fitness cost of synonymous recoding in bacteriophage ΦX174

**DOI:** 10.64898/2026.11.19.767875

**Authors:** James T. Van Leuven, Emma Altman, LuAnn Scott, Holly A. Wichman, Craig R. Miller

## Abstract

Synthetic biology allows for the synthesis of genomes from scratch. Although technically achievable, our abilities in designing genomes that result in live organisms are in their infancy. One important consideration is synonymous codon usage bias, which is present in most organisms and shaped by selection for efficient translation. Many studies have edited the synonymous codon usage patterns of viruses with the goal of viral attenuation, but these studies almost always build a single construct per design, so we know little about how repeatable the effects of recoding are. We designed and built 111 synonymously recoded ΦX174 genomes, applying four recoding strategies at up to five levels of attenuation to each of three genes, with three independently designed replicates of most combinations. Which gene we recoded mattered considerably more than how we recoded it. Recoding genes for the major capsid protein F, the major spike protein G and the DNA pilot protein H in full, with the least-used synonym codons from *E. coli* cost 12.0, 10.8 and 5.9 doublings per hour, respectively. Recoding strategy per se had no detectable effect once the number of codons changed was accounted for. Predicted mRNA folding stability also had no measurable effect on fitness when tested in gene F, whose wildtype transcript is highly stable and could easily be destabilized with synonymous codon rearrangements. While the number of codons changed was correlated with the degree of attenuation, divergence between independently designed replicates increased as more codons were edited. Replicates of designs with small fitness effects varied little, while replicates of strongly attenuating designs diverged by several doublings per hour. The fitness outcome of large-scale genome recoding is therefore substantially unpredictable from the design variables alone: even among independently designed genomes receiving the same treatment, fitness can diverge substantially.

## Introduction

Writing genomes is becoming routine, but designing them is not. Synthetic genomes have been built for viruses, bacteria, and yeast, yet our ability to predict the phenotype of a designed sequence remains limited. This gap matters most when the design intent is quantitative — that is, when the goal is a specific and stable reduction in replication rather than simply a viable organism.

Synonymous codon usage is one of the few genome-wide design variables that can be changed without altering any protein sequence. Codon usage is biased in nearly all organisms, the bias correlates with the abundance of cognate tRNAs, and it is strongest in highly expressed genes [1-5]. Synonymous sites are therefore not neutral. Changing them alters translation elongation rates, co-translational folding, mRNA stability, and protein output, sometimes by orders of magnitude [6-12]. A substantial part of this effect operates not through codons at all but through mRNA folding. In *Escherichia coli*, expression is better predicted by mRNA structure than by codon bias [7, 13, 14]. Reduced mRNA structure around the initiation site is therefore a conserved feature of genomes generally [15].

These observations have been put to practical use. Replacing preferred codons with rare ones, or disrupting preferred codon pairs, attenuates poliovirus [16], influenza virus [17], respiratory syncytial virus [18], chikungunya virus [19], and HIV-1 [20], among others (reviewed in [21]). Because attenuation is distributed across hundreds of sites, reversion is unlikely, which is the central appeal of the approach. Escape is not the same as reversion, however: a genome-scale deoptimized respiratory syncytial virus regained replication under selective pressure through a few mutations elsewhere in the genome rather than by reverting the recoded sites [22]. The mechanism of attenuation remains contested: the attenuating effect originally attributed to codon pair bias may instead reflect increases in CpG and UpA dinucleotide frequencies that accompany recoding [23]. Codon pair bias and dinucleotide composition covary under most recoding schemes, so distinguishing them requires designs built specifically to separate them.

Much of the work on synonymous recoding shares a limitation. For a given virus, gene, and recoding scheme, one sequence is typically designed and built. The enormous number of synonymous sequences encoding the same protein means that any single recoded genome is one draw from a large distribution of potential sequences. Where several synonymous recodings of a single gene have been built, the results have been sobering: Agashe et al. [24] recoded a key enzyme-coding gene of *Methylobacterium extorquens* entirely into both rare and frequently used synonymous codon, and found that both lowered expression and fitness. The fully codon-optimized version was the more costly of the two, reducing fitness by more than 90% against about 40% for the rare-codon version, which is the opposite of what codon-usage metrics predict; protein production tracked the frequency of anti-Shine– Dalgarno binding motifs rather than codon preference. Whether this is also true at the scale of a whole viral gene within a genome, and whether it depends on which gene is recoded, is unknown.

ΦX174 is well suited to answering these questions. It is a virulent bacteriophage of *E. coli* with a 5.4 kb single-stranded circular DNA genome. The virion consists of 60 subunits of the major capsid protein F, 60 subunits of the major spike protein G, and 12 subunits of the DNA pilot protein H [25]. Infection begins when a G pentamer contacts host lipopolysaccharide and is shed, after which F loops remodel to form a channel and H oligomerizes into a tube through which the genome is translocated [25, 26]. ΦX174 was the first genome to be assembled in its entirety from synthetic oligonucleotides [27], recoded variants can now be built by Golden Gate assembly [28], and we previously showed that deoptimizing whole ΦX174 genes lowers fitness, with the largest costs falling on the most highly expressed genes — so much so that the fully deoptimized G gene yielded no viable phage at all [29].

Here we ask how predictable synonymous recoding is. We designed four recoding strategies — replacing codons with the least preferred synonym, replacing them with a random synonym, shuffling the existing codons within the gene, and shuffling them to reach extreme predicted mRNA folding stabilities — applied each at up to five levels of recoding to each of genes F, G, and H, and built three independently designed replicates of most combinations. The resulting 111 genomes let us separate the contribution of the gene, the strategy, the extent of recoding, and the particular sequence drawn. We find that the gene matters a great deal, the strategy hardly at all, and the amount edited more than any of the deliberate design choices.

## Results

### Sequence space for ΦX174 recoding

We synonymously recoded genes for the major capsid protein F, the major spike protein G and the DNA pilot protein H in four ways. Codons that could be exchanged for a codon less frequently used in highly expressed *E. coli* genes were switched to the least-used synonym (“worst”). Codons were exchanged for a randomly chosen synonym (“random”). The existing codons of a gene were redistributed among the sites encoding the same amino acid, leaving the gene’s overall codon composition unchanged (“shuffle”). Finally, shuffled sequences were selected at the extremes of predicted mRNA folding stability (“mRNA folding”); this selection was performed in gene F only, whose wild-type transcript is unusually stable and could be destabilized with synonymous codon rearrangements (Results, Methods). Each strategy was applied at 25%, 50%, 75%, and 100% of the available sites, and the random strategy was additionally applied at a “max” level that changes every codon for which a synonym exists, including those that already use the least-preferred codon. This last treatment results in the most changed codons because some wildtype codons are already the least preferred and could not be changed in the 100% worst treatment.

The number of codons available to be edited differs substantially among the three genes and among strategies (Table 1). Gene F is 428 codons long and offers 296 sites that can be made worse; gene G is 176 codons with 108 that can be made worse; gene H is 329 codons with 197 that can be made worse. The shuffle strategy is the most constrained, because codons must be exchanged among synonymous sites within the gene, which leaves the gene’s codon adaptation index unchanged by construction and limited the shuffled constructs we built with at most 298, 97, and 188 changed codons for F, G, and H respectively. In every construct the first 21 nucleotides of the recoded gene were left unedited so that no design alters the sequence immediately around the start codon. Within each strategy and level, we selected sequence sets that were maximally divergent from one another, so that the three replicates of a treatment share as few of the same codons as possible (Methods).

**Table 1.**
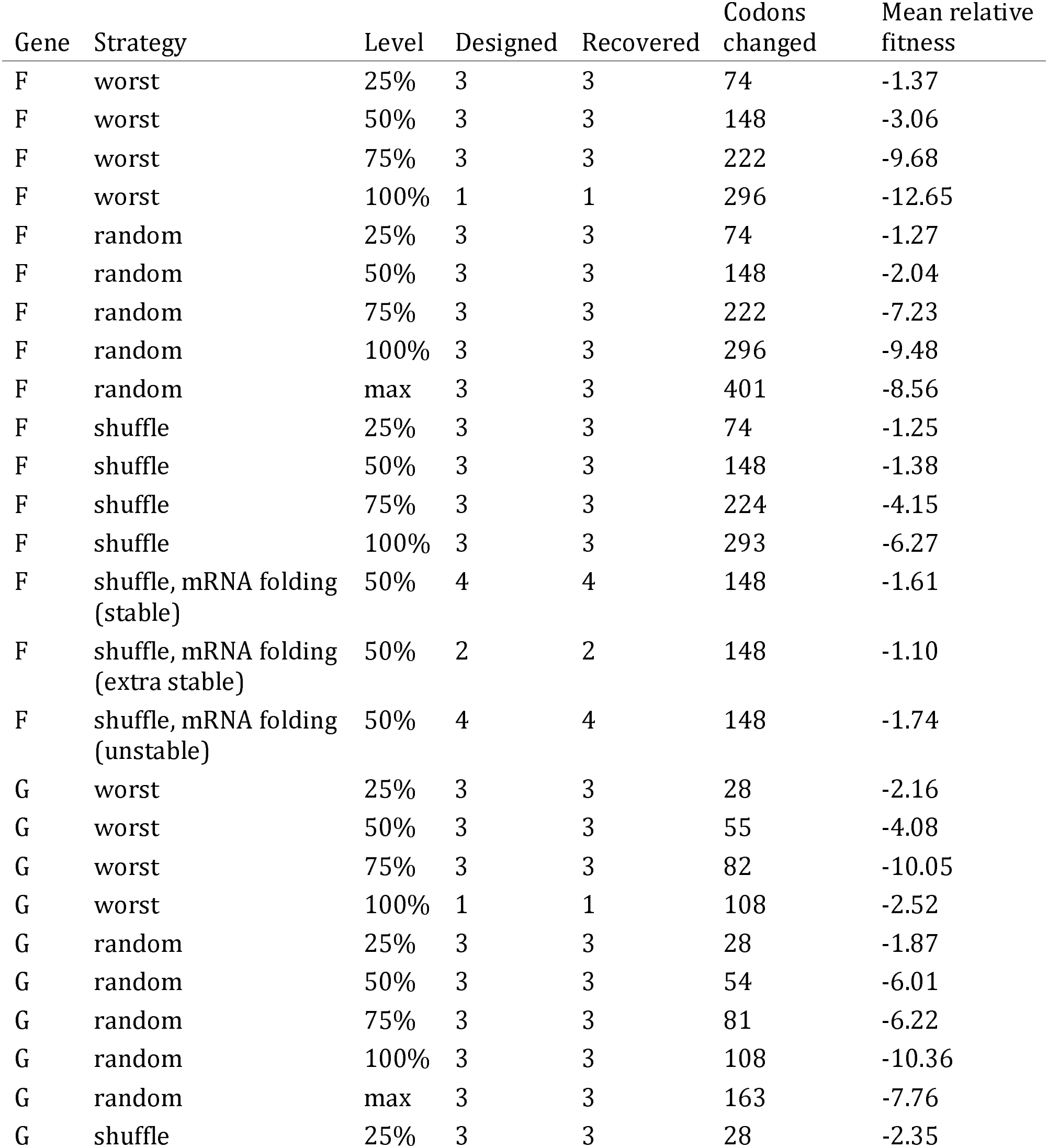
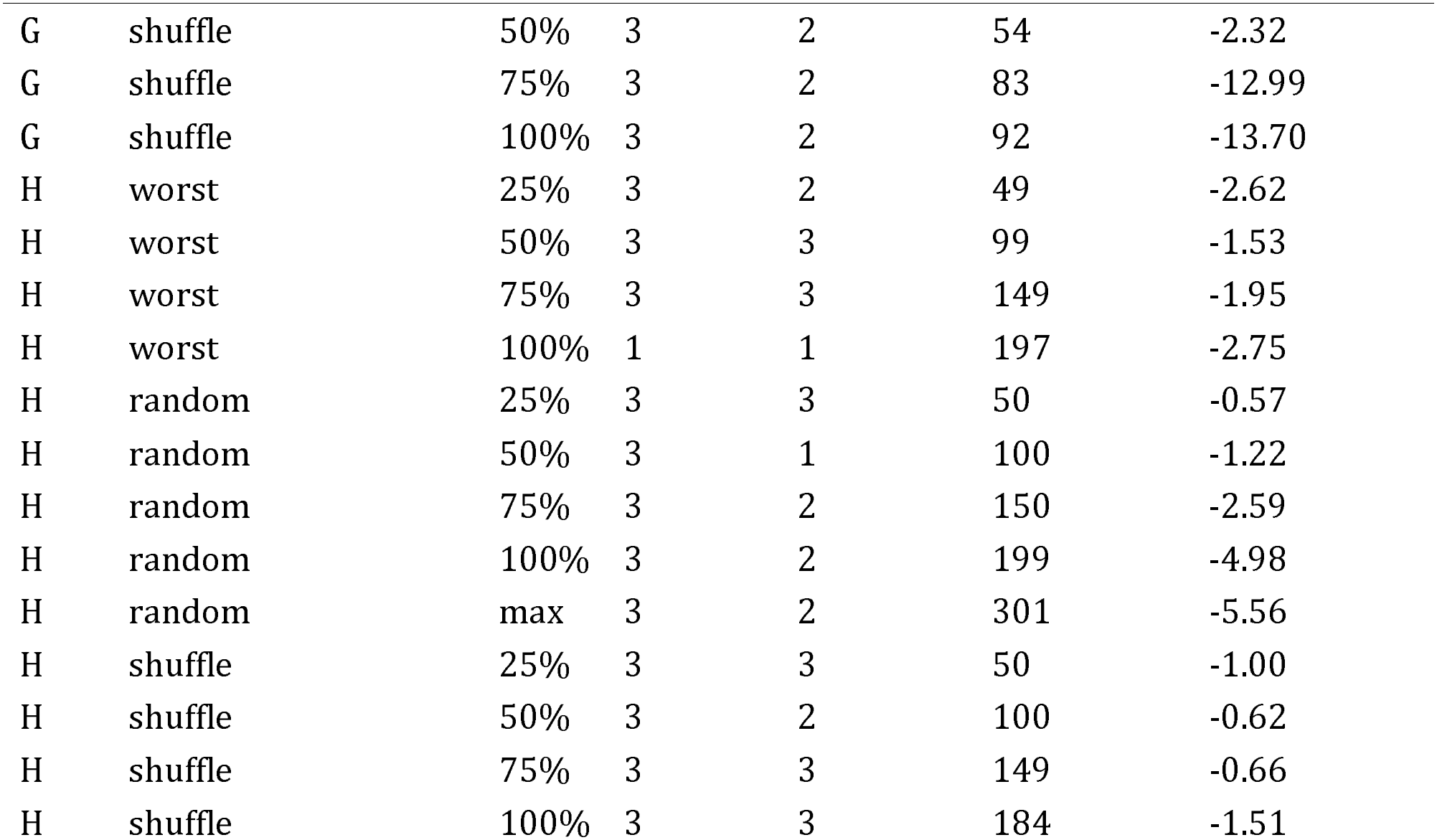
Recoding design and outcome. For each gene, strategy and level: the number of genomes designed, the number recovered and assayed, the number of codons changed within the recoded gene among recovered constructs, and their mean fitness relative to the same-day wild-type control (doublings per hour). Gene lengths are 428 (F), 176 (G) and 329 (H) codons. The “max” level was applied only to the random strategy.

Of the 121 genomes we attempted, 111 were recovered and fitness measured. Constructs were verified by whole-genome sequencing against their intended sequence (Methods). Comparing the intended sequences to the AF176027 reference confirms that no construct differs from the reference outside its targeted gene, and in every construct the recoded gene encodes the wild-type protein.

### Growth rate and competitive fitness agree

We measured fitness as a growth rate in liquid culture at low multiplicity of infection. To check that these values reflect competitive ability, we also competed constructs against one another in two independent pooled experiments and estimated fitness effects from changes in genotype frequency measured by PacBio sequencing. The two measures agree closely across the recoded constructs (Figure 1; Pearson r = 0.864, n = 98, P = 2.2 × 10^−30^ for the first competition experiment; r = 0.842, n = 99 for the second), and the two competition experiments agree with each other even more closely (r = 0.947). Rank agreement was similar (Spearman ρ = 0.89 and 0.90). We therefore use the growth-rate measurements throughout, as they are available for every construct.

**Figure 1.**
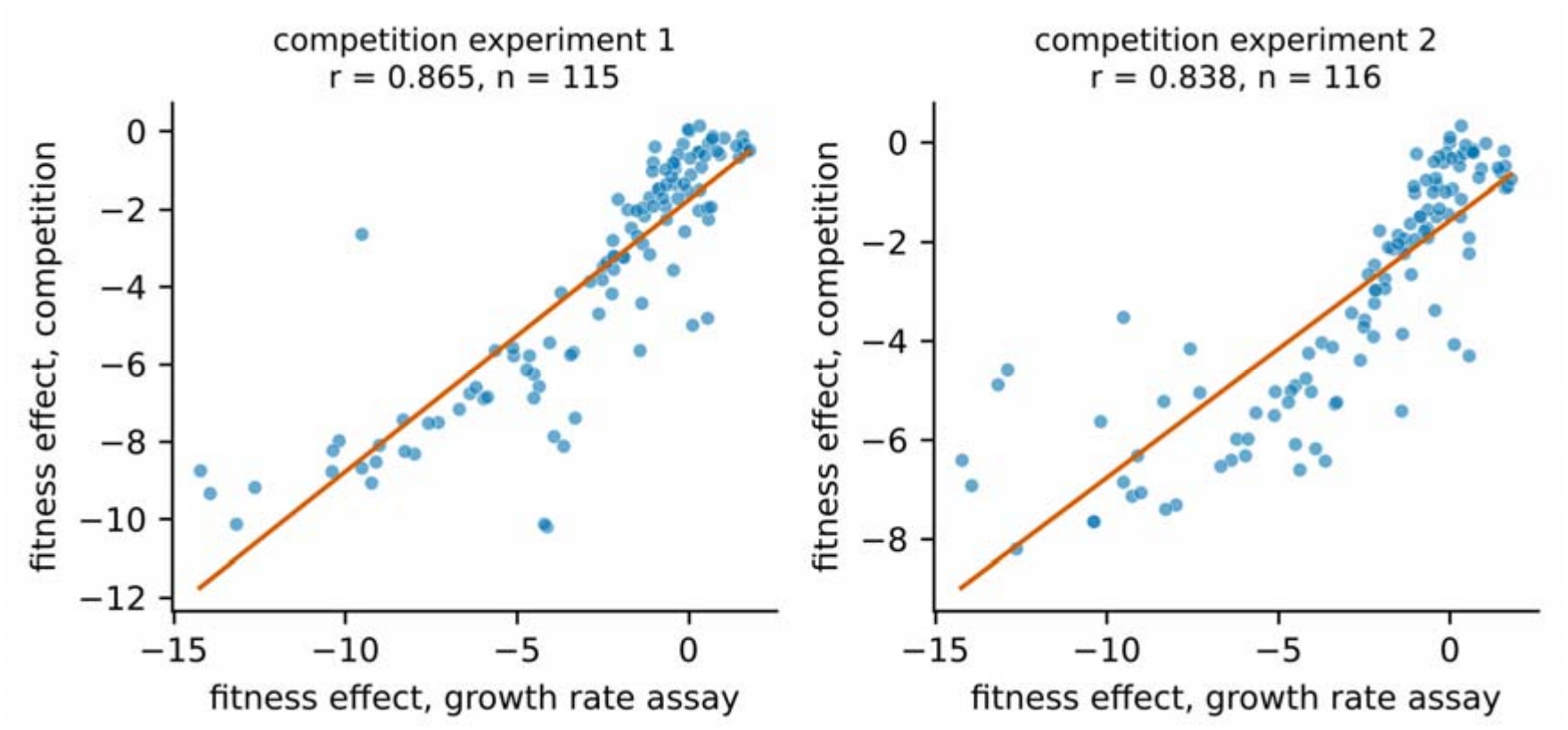
Growth-rate fitness compared with competitive fitness estimated from changes in genotype frequency during pooled competition, measured by PacBio sequencing. Each point is one construct; the two independent competition experiments are shown separately.

### Fitness declines with the extent of recoding at a rate that depends on the gene

Fitness fell as more codons were changed, in all three genes, but at markedly different rates (Figure 2, Figure 3, Table 2). Expressed as the cost of recoding an entire gene, the slope is −12.00 doublings per hour in gene F (95% CI −14.29 to −9.72, P = 8 × 10^−25^), −10.80 in gene G (−13.26 to −8.33, P = 8 × 10^−18^), and −5.94 in gene H (−8.67 to −3.20, P = 2 × 10^−5^). Allowing each gene its own slope improves the model substantially over a common slope (likelihood ratio 27.6 on 2 df, P = 1 × 10^−6^).

**Table 2.** Mean fitness relative to the same-day wild-type control (doublings per hour) by gene and extent of recoding, averaged over the worst, random and shuffle strategies, with the number of constructs in parentheses. Negative values indicate fitness below wild type. The mRNA folding constructs are omitted here because they exist at a single design level; they appear in Table 3.

| Gene | 25% | 50% | 75% | 100% | max |
|---|---|---|---|---|---|
| F | -1.30 (9) | -2.16 (9) | -7.02 (9) | -8.56 (7) | -8.56 (3) |
| G | -2.13 (9) | -4.36 (8) | -9.35 (8) | -10.16 (6) | -7.76 (3) |
| H | -1.24 (8) | -1.17 (6) | -1.62 (8) | -2.87 (6) | -5.56 (2) |

**Figure 2.**
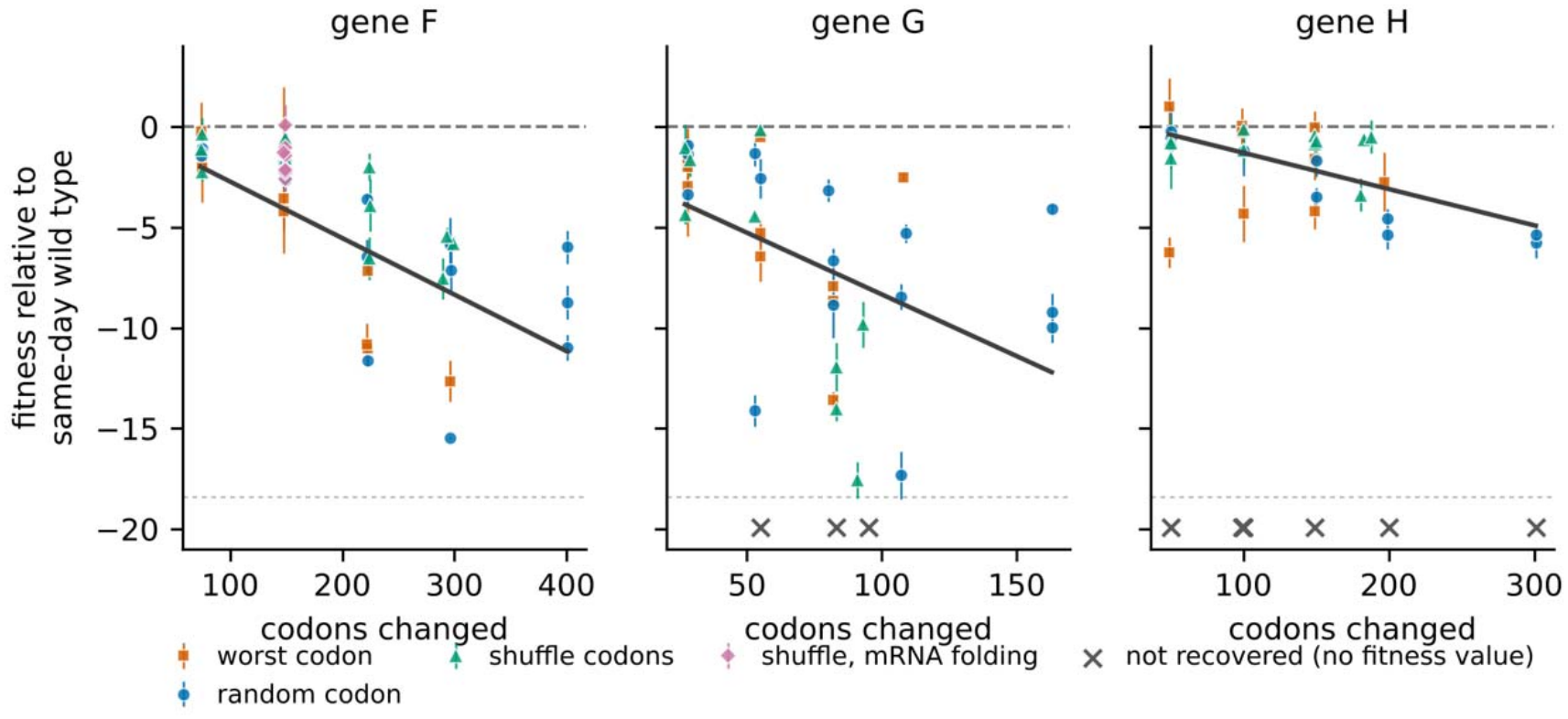
Fitness of recoded ΦX174 genomes, relative to the same-day wild-type control, against the number of codons changed. Each point is one construct, plotted as the mean of its assays; error bars are standard deviations across assays. Colors denote recoding strategy. The dashed line at zero marks the same-day wild-type level; the solid line is the per-gene slope from the mixed model described in the Methods, which is fitted to the worst, random and shuffle constructs and excludes the mRNA folding sets. Constructs are plotted at the number of codons they actually change. Crosses below the dotted rule mark the ten designs that could not be recovered, placed at the number of codons each was designed to change; these have no fitness value and lie outside the vertical scale. None are in gene F.

**Figure 3.**
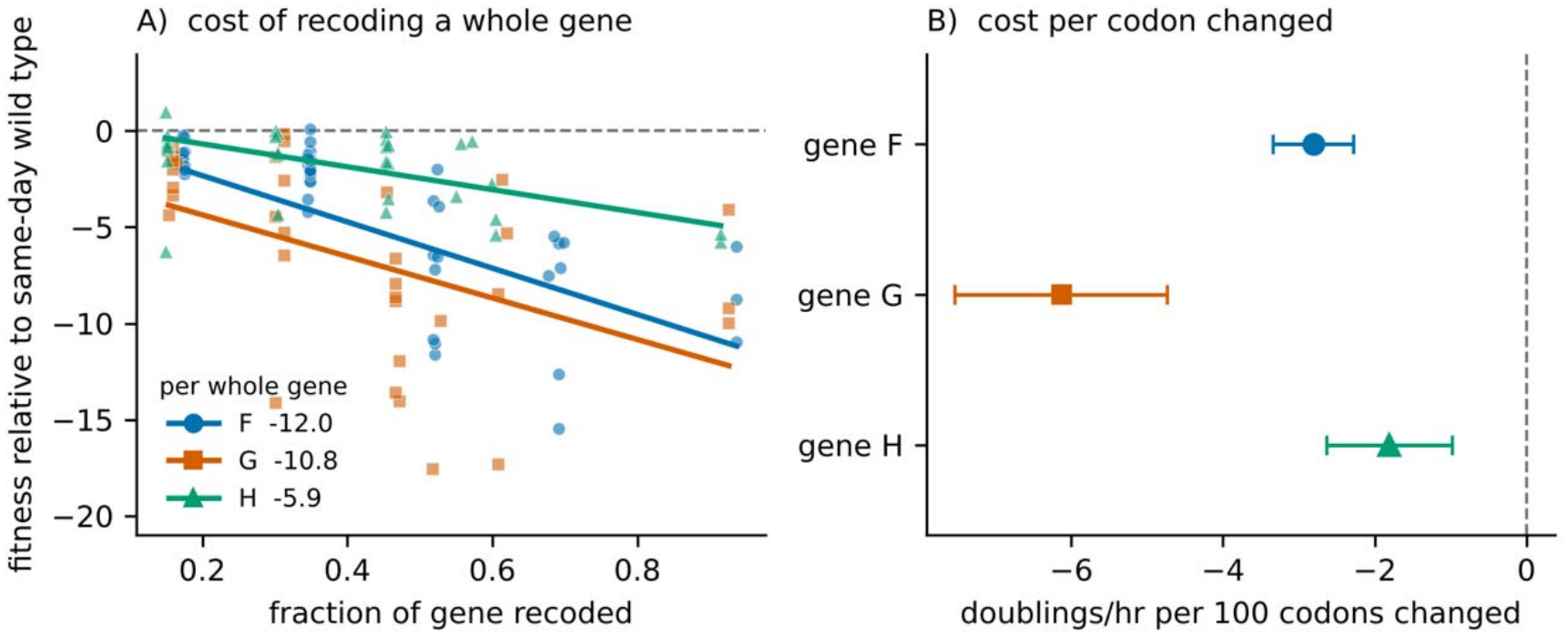
The cost of recoding depends on the gene. (A) Relative fitness against the fraction of each gene recoded; lines are the per-gene slopes from the mixed model described in the Methods, and the legend gives each slope in doublings per hour for a completely recoded gene. (B) The same model’s slopes expressed per 100 codons changed, with 95% confidence intervals. The two scalings give different orderings because the genes differ in length: gene G costs about as much as gene F to recode in full but more per codon changed, while gene H is the least sensitive on either scaling.

Because the three genes differ in length, the cost per gene and the cost per codon give different orderings. Per 100 codons changed, the slopes are −2.80 for gene F (95% CI −3.34 to −2.27), −6.13 for gene G (−7.53 to −4.73), and −1.80 for gene H (−2.63 to −0.97). The two scalings reorder the genes. Recoding a whole gene costs most in F, but gene G is less than half the length of F and costs almost as much, so per codon changed G is the most impacted of the three and F the second. That reordering is not a rounding artifact: the per-codon slopes of F and G differ by 3.33 doublings per 100 codons (SE 0.75, P = 9 × 10^−6^), and G differs from H by 4.33 (SE 0.82, P = 1 × 10^−7^). On either scaling gene H is the least sensitive, by roughly a factor of two per whole gene and a factor of one and a half to three per codon. The distribution of fitness effects at each recoding level is shown in Supplementary Figure 1.

### Recoding strategy has no detectable effect

Mean fitness by gene and strategy is given in Table 3. With the extent of recoding, gene identity, assay date and technician all in the model, recoding strategy does not improve the model. Replacing codons with the least-preferred synonym was indistinguishable from shuffling the gene’s existing codons (−0.01 doublings per hour, 95% CI −1.14 to +1.12, P = 0.99), as was random replacement (−0.57, 95% CI −1.39 to +0.24, P = 0.17). By contrast, the extent of recoding in the same model carries 12 doublings per hour across a fully recoded gene. This is a notable negative result. The “worst” and “shuffle” strategies are designed to do opposite things to codon adaptation: one drives the gene’s codon adaptation index as low as the genetic code allows, while the other leaves it exactly unchanged. That they are indistinguishable, with a confidence interval excluding costs larger than about one doubling per hour in either direction, indicates that the cost we observe is not attributable to codon preference but to the number of synonymous substitutions made.

**Table 3.** Mean relative fitness (doublings per hour) by gene and recoding strategy. These means are not directly comparable across strategies because the strategies differ in how many codons they change and because the mRNA folding sets exist at only one design level; the model described in the Methods controls for both. For gene F the mRNA folding column gives the stable and unstable groups separately; the two extra-stable gene F constructs are excluded there and averaged −1.10.

| Gene | worst | random | shuffle | shuffle, mRNA folding |
|---|---|---|---|---|
| F | -5.50 | -5.72 | -3.26 | -1.61 (stable), -1.74 (unstable) |
| G | -5.14 | -6.44 | -7.23 | — |
| H | -2.05 | -2.92 | -0.98 | — |

### Predicted mRNA folding stability

The transcript for Gene F is unusually structured, allowing us to test the effect of disrupting global mRNA stability. The wild-type gene F coding sequence has a Gibbs free energy (ΔG) of −393.9 kcal/mol, while 1,000 independently generated 50% shuffles of that same sequence, which by construction contain exactly the same codons in a different order, span −394.4 to −346.6 with a mean of −370.4 and a standard deviation of 7.6 (Figure 4A). The wild type lies 3.1 standard deviations below that mean. Wild-type codon order in gene F is therefore doing something to transcript structure that is not captured by codon composition alone, and if disrupting global structure carries a fitness cost anywhere in this genome, gene F is where it should appear.

**Figure 4.**
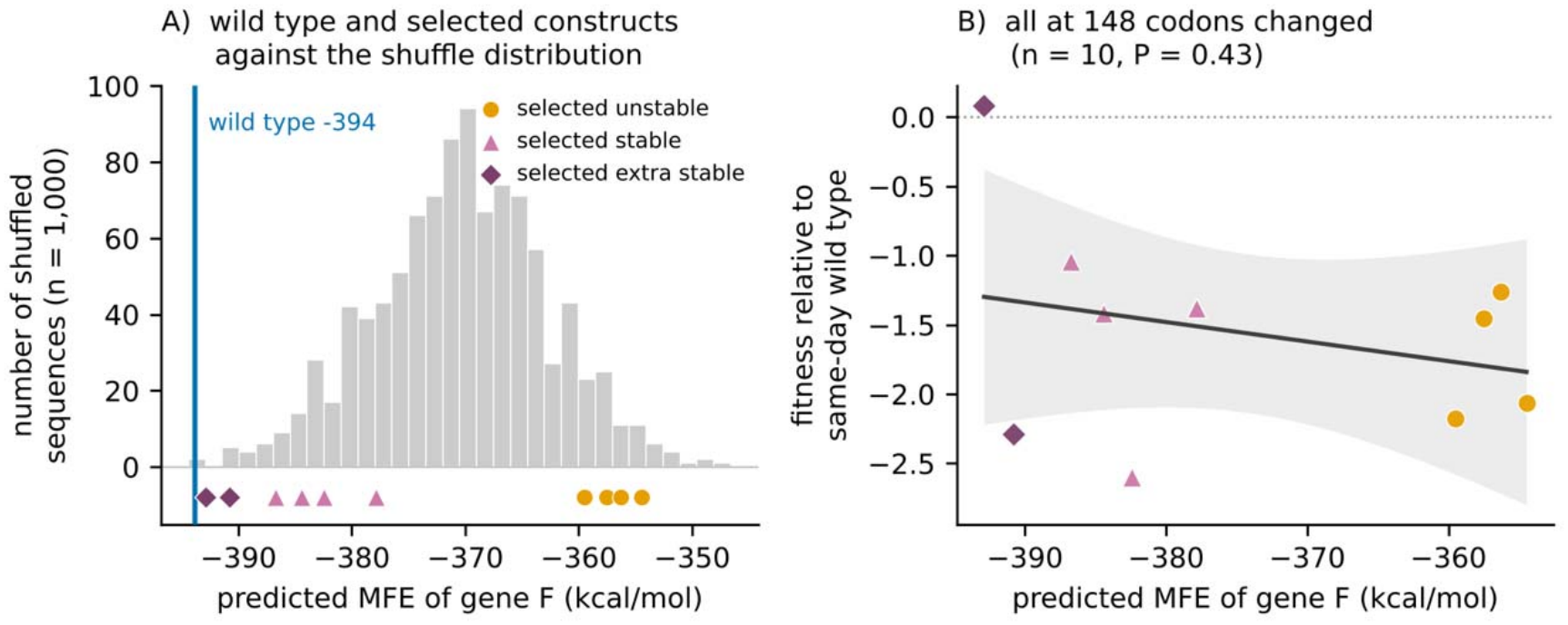
Predicted mRNA folding stability in gene F. All energies are ViennaRNA minimum free energies computed under the Turner 1999 parameters, so that the wild type, the candidate distribution and the built constructs lie on one scale. (A) The distribution of predicted minimum free energy across 1,000 independently generated 50% shuffles of gene F, which contain exactly the same codons as wild type in a different order. The blue line marks the wild-type value of −393.9 kcal/mol; one shuffle in the 1,000 is more stable. Triangles below the axis mark the ten constructs that were built — four selected from the stable tail, four from the unstable tail, and two selected to be more stable still — and are placed at their own energies rather than binned with the distribution. (B) Fitness relative to the same-day wild-type control against predicted folding energy for those ten constructs. All ten change 148 codons, half the available sites, and have codon composition identical to wild type, so recoding extent and composition are held constant and no covariate is required. The line is the ordinary least-squares fit with its 95% confidence band.

We built ten gene F sequences at the 50% level with codon composition identical to wild type: four drawn from the stable extreme of the candidate distribution, four from the unstable extreme, and two selected to be more stable still. The realised separation between the stable and unstable groups is 26.2 kcal/mol (−383.4, range −386.7 to −379.9, against −357.2, range −359.5 to −354.5; Welch P = 1.3 × 10^−5^). The extra-stable pair reach −390.8 and −392.9, approaching but not exceeding the wild-type value.

Fitness does not track this separation (Figure 4B). The stable group averages 1.61 doublings per hour below the same-day wild-type control and the unstable group 1.74, a difference of 0.13 (Welch P = 0.76), with the extra-stable pair 1.10 below, which differs from neither group (+0.57 against the other eight, Welch P = 0.71; this pair is only two constructs and the comparison is correspondingly weak). Because all ten constructs change the same number of codons and carry the same codons, fitness can be regressed on predicted folding energy directly, with no covariate required: the estimated effect across the 38.4 kcal/mol they span is −0.55 doublings per hour (95% CI −2.04 to +0.94, P = 0.42). Widening the test to all 22 shuffled gene F constructs, which span 41.0 kcal/mol and 74 to 298 codons changed, and controlling in the model for the number of codons changed, gives −0.27 doublings per hour across the full range (−2.34 to +1.80, P = 0.79). Restricting instead to the twelve shuffled constructs that were never selected on folding energy gives +0.77 across the 34.7 kcal/mol they span (−4.55 to +6.08, P = 0.75). The three estimates agree in that global mRNA folding is not predictive of fitness.

### Independently designed constructs with identical treatments diverge more than measurement error

The analyses above treat each treatment as a single quantity, but the three replicates within a treatment were independently designed sequences, and they often behaved very differently. The three fully recoded random-strategy G genomes, for example, grew 5.3, 8.5, and 17.3 doublings per hour slower than wild type. These constructs were designed identically, each changing 108 of gene G’s codons to randomly chosen synonyms, yet they span 12 doublings per hour: one is moderately attenuated and another barely grows at all (Figure 5A). Adding construct identity to the model as a random effect nested within treatment separates the variation among independently designed constructs from the variation among repeated assays of the same construct. The among-construct standard deviation is 2.61 doublings per hour against an assay-to-assay standard deviation of 0.96 (Figure 5B), so which particular sequence was built accounts for 88% of the variation that recoding strategy does not explain, and constructs sharing a treatment differ from one another 2.7 times as much as repeated measurements of the same construct.

**Figure 5.**
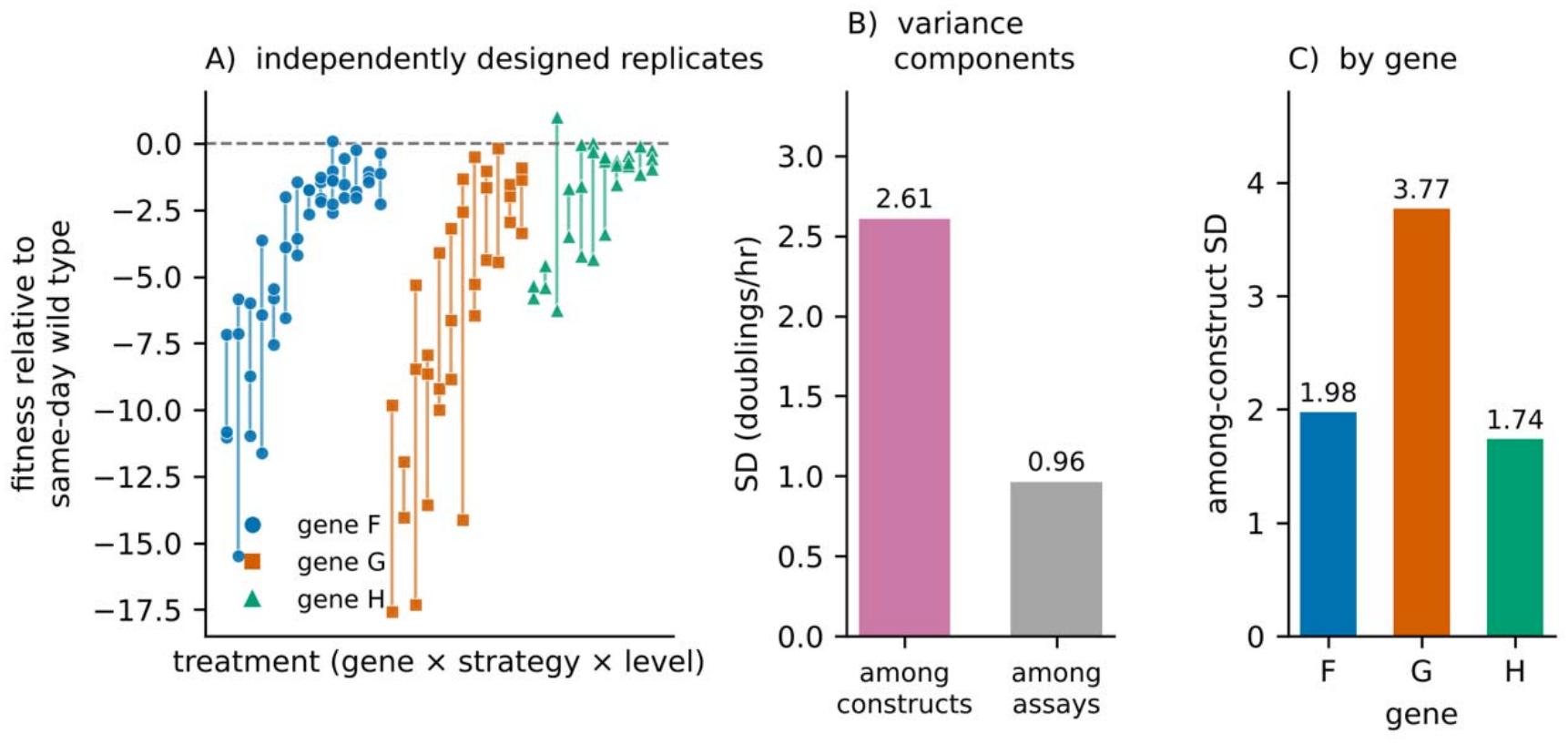
Divergence among independently designed replicates. (A) Relative fitness of the constructs within each treatment, with a line connecting the extremes; constructs sharing a design often differ widely. All treatments with at least two recovered constructs are shown, ordered by gene and then by mean fitness. (B) Variance components: the standard deviation among constructs within a treatment against the standard deviation among repeated assays of the same construct. (C) The same among-construct standard deviation fitted separately within each gene. Panels B and C use all 111 constructs.

Among-replicate standard deviation correlates with treatment mean fitness (r = −0.588, P = 1 × 10^−4^, across 37 treatments with at least two recovered constructs). Recoding designs that barely attenuate are reasonably reproducible. Designs that attenuate substantially are not, and the loss of reproducibility sits in the range that an attenuation study would actually target. Fitting the variance components separately to the two halves of that range, among treatments whose constructs average within 5 doublings per hour of wild type the among-construct standard deviation is 1.49 against an assay standard deviation of 1.03; among treatments averaging more than 5 doublings below wild type it is 4.02 against 0.79. The divergence is also gene dependent. Fitting the variance components within each gene, the among-construct standard deviation is 3.77 doublings per hour in gene G, 1.98 in gene F, and 1.74 in gene H (Figure 5C), tracking the degree of attenuation each gene sustains.

### Construction failures

Ten of the 121 attempted genomes could not be recovered after three independent ligation and transformation attempts. Failures were strongly structured by gene: none of 47 gene F constructs failed, against 3 of 37 for gene G and 7 of 37 for gene H (Fisher’s exact test, F against H, P = 0.0023). They were not concentrated at the highest recoding levels: four occurred at the 50% level, two each at 75% and 100%, and one each at 25% and max.

This pattern bears on the interpretation of gene H’s shallow fitness slope. The gene whose recoded genomes are least affected when they can be measured is also the gene whose genomes most often could not be built at all, and an unrecoverable genome plausibly represents a severe defect rather than a missing observation. Taken together, the two observations suggest that recoding gene H is usually close to neutral but occasionally catastrophic, which is a different claim from uniform tolerance and one that is consistent with the general unpredictability we document above. We report fitness analyses with these constructs excluded, and note that this exclusion is not random with respect to gene.

## Discussion

We set out to determine the predictability of virus attenuation using synonymous recoding. Of the three things we deliberately varied — which gene, how much of it, and by what strategy — only the first two mattered, and together they still left fitness variance largely unexplained. Thus, we concluded that there is a high amount of unpredictability when making many synonymous changes to genes.

### Which gene is recoded matters more than how

For all genes that were recoded fitness decreased proportional to the number of changes made, but at rates differing by about a factor of two per whole gene: 12.0 doublings per hour for gene F, 10.8 for gene G, and 5.9 for gene H. Considering the effects as a function of the proportion of the gene edited changed the gene rankings, with gene G steepest at −6.13 doublings per 100 codons, gene F at −2.80, and gene H shallowest at −1.80. On either scaling gene H is the least sensitive. All three targets are structural genes, which is the class in which translational selection on phage codon usage is most evident: across 74 phages, head and tail proteins are significantly more biased toward host-preferred codons than non-structural proteins [30]. The differences we see are therefore differences within the gene class expected to be most strongly optimized, not between optimized and unoptimized genes. Protein abundance is an obvious explanation for fitness impacts. The ΦX174 virion contains 60 copies each of F and G [31] but only about 12 of H, which forms a DNA-translocating tube rather than part of the capsid shell [25]. Selection on codon usage scales with translational flux, so the codon usage of highly expressed genes is both more strongly optimized and more costly to disrupt [1-3, 32]. The interpretation is consistent with our earlier whole-gene deoptimization of ΦX174, in which recoding the highly expressed genes J, F and G produced larger fitness decreases than recoding the lowly expressed A and H, and the fully deoptimized G gene could not be recovered at all [29]. Transcript abundance follows the same ranking. RNA sequencing of ⍰X174 infection places gene F at roughly twice the transcript level of gene G, and gene G about half again above gene H [33], which is the order in which recoding a whole gene impacts fitness. The correspondence fails on the other scaling. Per codon changed, gene G is the most expensive of the three and differs from gene F by 3.33 doublings per 100 codons (SE 0.75, P = 9 × 10^−6^), despite sitting between F and H in both transcript abundance and virion copy number. Abundance therefore tracks the cost of recoding an entire gene, largely because the most abundant gene is also the longest, and fails to explain what an individual substitution costs. With only three genes tested, we offer this interpretation as a consistency rather than a likely cause.

A related caution concerns the interpretation of gene H’s shallow slope. H is the least sensitive gene among the genomes we could assay, but it is also the gene whose genomes most often could not be assembled at all: 7 of 37 H constructs failed compared to none of 47 F constructs. If unrecoverable genomes represent severe fitness defects, as seems likely, then the distribution of outcomes for recoded H is better described as usually near-neutral but occasionally catastrophic than as uniformly tolerant. That description is itself a statement about unpredictability, and it fits the rest of our results.

### Recoding strategy did not matter

We expected the “worst” strategy to be the most damaging, since it drives codon adaptation as low as the genetic code permits, and the “shuffle” strategy to be the least, since it leaves the gene’s codon composition unchanged. The two were indistinguishable, with a confidence interval excluding a difference larger than about one doubling per hour in either direction, while the extent of recoding in the same model carried twelve doublings per hour across a fully recoded gene. Because two technicians ran almost disjoint sets of strategies, we estimated the technician difference from the wild-type assays, which both technicians performed throughout the experiment; that difference is small and the null result does not depend on it (−0.47 doublings per hour, 95% CI −1.42 to +0.48, see Methods).

Many interpretations of these results are possible. The first is that the cost of recoding in this system is driven mostly by some factor not accounted for in our recoding strategy and the likelihood of changing this unaccounted for variable increases as the number of edited sites increases. However, we calculated many properties such as codon pair bias and GC content of the sequences that we designed (see [29] for a complete description of sequence properties) and none offer improved predictive power. Another possibility is specific to this phage. 11X174 begins further from its host’s codon preferences than most phages do: among 49 coliphage genomes, the Microviridae show the weakest correlation between phage and *E. coli* codon usage of any family examined, with a mean rRSCU of 0.32 against 0.42 to 0.70 for the double-stranded DNA families, a difference attributed to the high C11T mutation bias of single-stranded genomes [34]. A genome that is already poorly adapted has less to lose, which would compress the distance between driving codon usage further from the host optimum and merely rearranging it. These readings are not mutually exclusive, and the mechanistic ambiguity resembles the reinterpretation of codon pair bias effects as dinucleotide composition effects [23]: metrics that are easy to compute and easy to manipulate need not be the metrics that matter. Codon order is itself known to carry information: synonymous codons that reuse the same tRNA are clustered rather than randomly placed, and in *Saccharomyces cerevisiae* anticorrelated sequences are translated more slowly than their correlated synonymous counterparts [35]. Our shuffle constructs hold codon composition fixed while rearranging codon order, and so disrupt both dinucleotide content and any such autocorrelation, yet they attenuate in proportion to the number of sites moved rather than to any property of the rearrangement itself. That makes this data set a good one for testing order- and composition-based explanations directly in future work.

For the practical goal of attenuation, the irrelevance of strategy is convenient. If fitness reduction depends mainly on how many synonymous sites are changed and not on how they are changed, then attenuation can be tuned by the number of edits, and the designer is free to use the remaining degrees of freedom for other purposes, such as avoiding restriction sites, maintaining assembly overhangs, or minimizing the chance of recombination with circulating strains.

### mRNA folding stability

Folding free energy is among the better predictors of expression in *E. coli*, though the published relationships concern local structure near the translation start rather than the stability of a transcript as a whole [7, 13, 14]. Our designs deliberately left the first 21 nucleotides of each gene unedited so as not to disturb translation initiation, and the manipulation is consequently a global one. Ten gene F constructs, each changing 148 codons and carrying exactly the same codon composition as wild type, span 38 kcal/mol of predicted folding energy; the fitness difference across that span is bounded between −2.0 and +0.9 doublings per hour. Widening the comparison to all 22 shuffled gene F constructs and 41 kcal/mol does not change the answer. Thus, in ΦX174 at least, changing global mRNA stability seems to have no impact on fitness. Whether local structure at the initiation codon would behave differently is a separate question that our design did not address, since we held that region constant on purpose. This hypothesis is one we would encourage others to test directly by selecting on the folding energy of a window around the start codon.

### Unpredictability and what it means for genome design

The result we find most consequential concerns among-construct variation. Independently designed genomes receiving identical treatments differed from one another with a standard deviation of 2.61 doublings per hour, against 0.96 for repeated assays of the same construct, so construct identity accounted for 88% of the variation that treatment (what gene and % recoded) did not explain. In the starkest case, three identically designed G genomes spanned 12 doublings per hour. We should be clear about what this does and does not suggest. The variation that we observe is biological rather than methodological as it greatly exceeds assay variability, and it is not a batch effect: with assay date modelled explicitly as a variance component, the date term has a standard deviation of 0.56 doublings per hour against 2.60 for construct identity.

This among-construct variance is strongly conditional on how much the treatment attenuates. Among treatments whose constructs average within 5 doublings per hour of wild type, replicates diverge with a standard deviation of 1.49, not far above the 1.03 of the assay itself. Among treatments averaging more than 5 doublings below wild type, that figure is 4.02 (r = −0.588 between among-replicate standard deviation and treatment mean, P = 1 × 10^−4^). The same pattern appears across genes: gene G, which sustains the largest fitness losses per codon, has an among-construct standard deviation of 3.77, while gene H, which is barely attenuated, has 1.74. Recoding is therefore reproducible while it remains nearly neutral and becomes unpredictable over exactly the range of attenuation that an attenuated-vaccine design would target.

What we cannot yet say is where the variation comes from. Epistasis among synonymous sites is a natural explanation, and our earlier observation that deoptimized ΦX174 genes combine non-additively [29] points the same way, as does the general finding that synonymous recoding of a single gene produces idiosyncratic, metric-defying effects [24]. We had hoped to distinguish the identity of the substituted codons from their position by exploiting the fact that replicates at high recoding levels are forced to edit overlapping sets of sites. However, at the fully recoded levels replicates share about three-quarters of their edited positions, so position and identity remain confounded. Separating them will require a design in which the edited positions are held fixed and only the choice of synonym varies.

### Limitations

Our fitness measure is a growth rate in a single host under a single laboratory condition, and although it agrees closely with competitive fitness measured by sequencing (r = 0.864 and 0.842 in the two independent competition experiments), both were measured in the same environment. We recoded three genes of one phage, so the abundance explanation for gene-specific sensitivity rests on a comparison of three points and, as noted, does not account for the difference between F and G. Ten constructs could not be built, and because failures were concentrated in genes G and H, the surviving sample is not random with respect to gene; this censoring works against the tolerance we report for gene H rather than for it.

## Tables

Some codons in each wild-type gene already use the least-preferred synonym and cannot be made worse, which caps the “worst” strategy at 296, 108 and 197 codons for F, G and H. The random strategy at the max level changes every codon for which any synonym exists: 401, 163 and 301 codons. Shuffling is more constrained still, because codons must be exchanged among synonymous sites within the gene; the shuffled constructs we built reach at most 298, 97 and 188 changed codons. Codon and nucleotide changes are counted against our laboratory ancestor AF176027. The first 21 nucleotides of each gene were excluded from recoding in every design.

†Constructs selected on predicted mRNA folding energy in gene F. Three groups are reported: four stable, four unstable, and two extra-stable, each changing 148 codons with codon composition identical to wild type.

## Figures

**Supplementary Figure 1.**
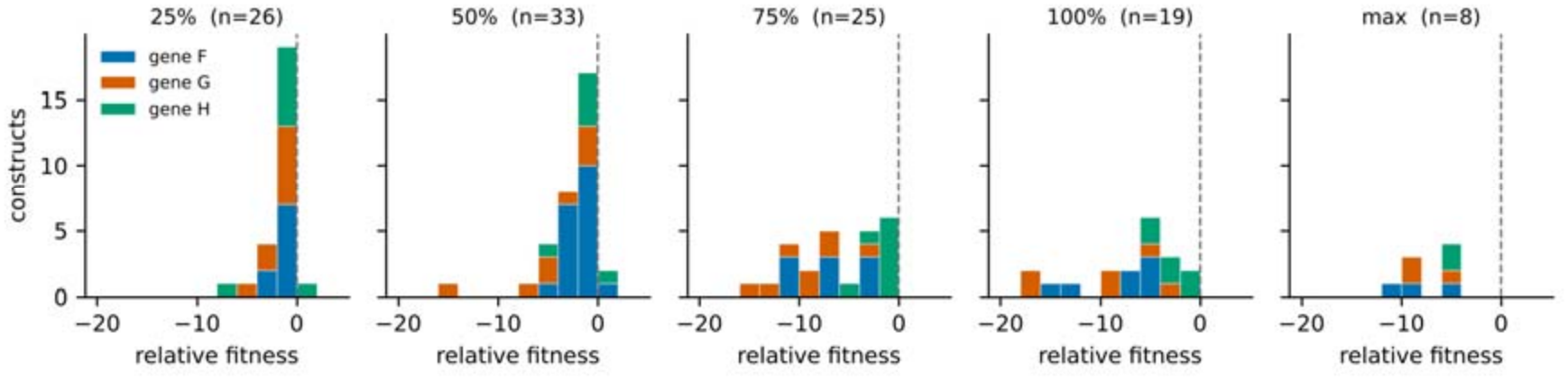
Distribution of fitness effects by recoding extent. Relative fitness of all 111 constructs, binned by nominal recoding level and coloured by gene. The distribution broadens and shifts downward with recoding extent, and gene H contributes most of the near-neutral constructs at every level.

## Methods

### Bacteriophage and host strains

A laboratory strain of bacteriophage ΦX174 (GenBank accession number AF176034) was used in this study. All experiments were carried out using *E. coli* C as a host in modified Luria–Bertani medium (10 g/l tryptone, 5 g/l Bacto yeast extract, 10 g/l NaCl, 2 mM CaCl_2_).

### Recoding algorithm

To generate synonymously recoded gene sequences, we created an algorithm that searches sequence space and outputs a given number of unique gene candidates. The algorithm starts by determining all possible synonymous mutations and their associated changes in codon preference from *E. coli* codon preference values [5]. For each level of recoding (25%, 50%, 75%, 100%, and max), the algorithm changes available codons until the desired level is achieved, using three strategies. For the “worst” strategy, the codon is exchanged for the least preferred synonym. The “random” strategy picks a synonymous codon at random. Because not all codons can be switched for worse codons, some already being the least preferred, we restricted the number of mutations made to any given construct to the total number of codons that can decrease in codon preference. This keeps the total number of mutations consistent across recoding strategies. The “max” level is the exception to this rule and was applied only with the random strategy, which does not require that a worse codon be available. We also attempted the “max” level with the shuffle strategy but could not edit more codons than were edited at the 100% level, because shuffling requires swapping codons within a gene and so leaves the overall codon adaptation index unchanged. Codon composition is preserved exactly by the shuffle strategy in genes F and H, which we verified across all 1,000 candidates in each gene. In gene G every shuffled candidate differs from wild type by a single asparagine codon, carrying eight AAT and three AAC where wild type has seven and four; the eleven asparagine sites are otherwise redistributed as intended. The effect on codon adaptation is negligible, but gene G shuffles are not strictly composition-preserving and we note it where the distinction matters. We excluded codons required to generate the sticky ends used for ligating fragments together, and we excluded the first 21 nucleotides of each gene so that no design would alter the sequence immediately around the start codon, where mRNA structure is known to affect translation initiation. The earliest edit in any construct therefore falls at the seventh codon. For each level and recoding strategy, we generated 1,000 candidate sequences.

### Selection of divergent sequence sets

To increase the independence of our observations, we selected sequences that were maximally divergent from one another within a given strategy. For each strategy (random, worst, and shuffle) we took the following steps. (1) For each level of recoding (25%, 50%, 75%, and 100%), we sampled n = 1,000 sets of three sequences (trios). (2) For each trio we calculated the three pairwise nucleotide sequence differences and took the minimum. We then identified which trio among the 1,000 had the largest minimum pairwise difference. (3) We repeated steps 1 and 2 ten times to obtain ten sequence trios with high within-trio divergence. (4) After doing this for each level within a strategy, we examined every possible way to combine one trio from each level (10 from 25% × 10 from 50% × 10 from 75% × 10 from 100% = 10^4^ combinations). For each combination we performed all pairwise sequence comparisons among the 12 sequences involved (4 levels × 3 sequences per level) and scored the combination as the minimum value. (5) We identified the combination with the largest minimum score and this defined the set of trios used in our experiments. We did not control for divergence between strategies.

There were two minor exceptions to this procedure. For the random strategy there was a fifth level of recoding (“max”), so there were 10^5^ combinations to search across. For the worst strategy the 100% level was a single sequence, that containing the worst codon at every mutable site, so there was variation at only three levels and 10^3^ combinations to search across; we nevertheless included the single 100% sequence when searching for the best combination across levels in step 5.

To identify divergent sequences with contrasting effects on predicted transcript stability, we focused on the 50% shuffle level and sampled at both extremes. We calculated the folding stability of all 1,000 candidate 50% shuffle sequences per gene using mfold 3.6 [36], sorted them by predicted folding free energy, and identified the top and bottom 5% (n = 50 each). We then repeated steps 1 to 5 above with the top and bottom 5% defining our groups, sampling sets of four (quartets) rather than three. In gene F, whose wild-type transcript is more stable than essentially all of its synonymous rearrangements (Results), nearly all shuffled sequences were less stable than wild type; a very small number were more stable, and without regard to sequence divergence we selected the two most stable of these and appended them to our dataset. Sequence selection was carried out in R with the aid of the Biostrings package. The same selection was attempted in genes G and H, but the chosen indices were applied to the 100%-shuffle candidate pool rather than to the 50% pool from which the folding energies had been computed. The sequences built for those two genes are therefore 100% shuffles whose predicted folding energies bear no relation to the intended stable and unstable groups. We exclude all 15 recovered gene G and gene H constructs from this study, and they are not counted among the 111 genomes analysed here; the mRNA folding contrast is reported for gene F only.

### Synthetic ΦX174 genomes

A phage assembly platform for ΦX174 was devised following Faber et al. [28]. The ΦX174 chromosome was divided into 14 genomic fragments designed to avoid host cell toxicity by separating genes from their promoters and breaking large genes into multiple segments. Each segment is flanked by unique four-nucleotide overlaps of wild-type ΦX174 sequence so that they could be cut from the cloning vector using BsmBI and ligated. Fragments were synthesized as commercial oligonucleotide pools and assembled by Golden Gate ligation as described in Faber et al. [28]. The ligation mix was transformed by electroporation into 100 μl competent *E. coli* C cells. The transformation mix was resuspended with 1 ml of ΦLB and plated immediately. The ΦLB was added to 3 ml of ΦLB top agar and plated onto a ΦLB agar plate. After four to five hours of incubation at 37 °C, recombinant phage plaques were visible and plates were removed from the incubator. Plaques were picked into ΦLB with 50 μl of chloroform. These stocks were used for sequencing and fitness assays. To verify that each recombinant phage encoded the intended sequence, the resulting phage genome was sequenced in its entirety as previously described [37]. Constructs that could not be recovered on the first ligation attempt were attempted two additional times; constructs that failed all three attempts were recorded as construction failures and excluded from fitness analyses.

### Sequence verification

Each construct’s intended full-length genome sequence was compared to the AF176027 reference across all 5,386 positions, and we recorded the positions and identities of all differences. Intended sequences were available for 102 of the 111 recovered constructs. The nine exceptions, all gene G shuffles, were reconstructed from the design records: the candidate sequence pool and the recorded index of the selected sequence within it. We validated this reconstruction procedure on the one gene G shuffle whose intended sequence is present in our sequence collection (the 100% shuffle replicate 3, which did not itself yield a virus): the reconstruction reproduced it exactly, confirming both the indexing convention and the mapping of selection slots to replicate numbers. For each construct we recorded the positions and identities of all differences, confirmed that all differences other than three fixed differences of our laboratory ancestor (positions 1650, 4518, and 4784) fell within the targeted gene, and confirmed that the translated product of the recoded gene was identical to wild type. Gene boundaries were taken as F 1001–2284, G 2395–2922, and H 2931–3917.

### Measuring viral fitness

Fitness assays and fitness calculations were performed as previously described [37]. The assay is a determination of growth rate at low multiplicity of infection in 10 ml ΦLB, carried out at 37 °C. Host cells were prepared by growing to approximately 10^8^ cells/ml and aliquoted into 8.5 ml of warm LB just prior to adding phage. Fitness is expressed as the number of doublings per hour, the log_2_ of the ratio of phage concentration at the end of the assay to phage concentration at time zero, divided by the elapsed time in hours. In the normalised data set analysed here, one construct was measured four times, 123 three times and two twice. Assays were performed by two people; as reported in the Results, the two each ran an almost disjoint set of recoding strategies, though genes were divided between them in nearly equal numbers. Virus titers were determined on ΦLB agar plates with 0.7% top agar. Assays sampled at 40 min and the log_2_ increase was multiplied by 1.5 to express fitness per hour, as in Wichman et al. [37].

Wild-type ΦX174 was assayed on 62 dates, providing a same-day control on 52 of the 53 dates on which constructs were assayed and for 377 of the 386 construct assays. All analyses are performed on relative fitness, defined as a construct’s fitness minus the mean fitness of the wild-type assays performed on the same date, in doublings per hour. The nine assays without a same-day control are excluded from analyses of relative fitness; all 111 constructs retain at least two normalisable assays.

Two technicians performed the fitness assays over the course of about one year. Wild type was always measured alongside recoded constructs. Therefore, we analyzed raw fitness in a single mixed model that includes the wild-type assays, with assay date as a random effect and technician as a fixed effect. Because wild type was measured by both technicians and is unrelated to recoding strategy, it identifies the technician term independently of the construct comparisons, which a model fitted to constructs alone cannot do. All inferential statements below come from this model (310 construct assays and 73 wild-type assays across 63 dates); the tables report descriptive means relative to the same-day wild-type control. The technician term is small and not distinguishable from zero: −0.47 doublings per hour (95% CI −1.42 to +0.48, P = 0.33), consistent with the −0.32 estimated from the wild-type assays alone.

### Competition assays and sequencing-based fitness estimates

Constructs were pooled and competed against one another in two independent experiments, each with three replicate cultures. Samples were taken from the initial pool and after 40 and 80 minutes of growth. Amplicons spanning the genome were sequenced on the PacBio platform. Reads were quality-filtered and length-filtered to 3,000–3,500 bases with fastp, aligned to a reference set containing all 127 genotypes with bwa mem, filtered to mapping quality ≥ 20 and to fewer than 20 mismatches per read, and counted per genotype with seqkit. Genotype frequencies were computed as the proportion of assigned reads in each sample, and fitness effects were estimated from the change in genotype frequency over time by regression.

### mRNA folding stability

Predicted minimum free energies of recoded coding sequences were computed with RNAfold from the ViennaRNA package version 2.7.2 [38], folding the complete coding sequence of the recoded gene. Ensemble free energies were computed from the partition function for comparison. Sequence selection at design time used mfold 3.6 [36], which implements the Turner 1999 nearest-neighbor parameters [39], whereas ViennaRNA 2.7 defaults to the Turner 2004 set. We therefore report folding energies under both parameter sets. On the 22 ordered gene F sequences for which design-time mfold values are available, the Turner 1999 parameters reproduce mfold with r = 0.983 and a mean offset of −16.2 kcal/mol (SD 2.9), against r = 0.954 and +11.7 kcal/mol (SD 4.4) for the Turner 2004 defaults. Those 22 sequences were selected at the extremes of the folding distribution, however, and the agreement across the unselected candidate pool is appreciably weaker: on all 1,000 gene F candidates, mfold and the Turner 1999 implementation correlate at r = 0.931 with a residual standard deviation of 2.79 kcal/mol about the fitted relationship. The two therefore rank sequences similarly but not interchangeably, and absolute values are not comparable between tools at all. The gene F folding analysis reported in the Results uses ViennaRNA values computed under the Turner 1999 parameters throughout, so that the constructs, the candidate pool and the wild-type sequence are all placed on one scale; the design-time mfold values exist only for the 22 ordered sequences and not for the wild type or the full pool. All comparisons are made within a gene under a single parameter set.

To ask whether a wild-type coding sequence is unusually structured for its codon composition, we folded the wild-type sequence of each gene and compared it with that gene’s 50%-shuffle candidate pool, which by construction contains the same codons in a different order. For gene F all 1,000 candidates were folded. We report the number of shuffles more stable than wild type and the position of the wild type in standard deviations from the shuffle mean. No mfold value was computed for any wild-type sequence at design time, so this comparison uses ViennaRNA under the Turner 1999 parameters for the wild type and the candidate pool alike, which is the only way to place the two on a common scale. The candidate pools available to us were regenerated after the constructs were ordered and are not the particular draws from which the built sequences were selected; they serve here as reference distributions of synonymous rearrangements of each wild-type gene, which is what this comparison requires.

### Statistical analysis

Analyses were performed in Python using pandas, SciPy and statsmodels. Fitness was analyzed on its measured scale (doublings per hour) in a linear mixed model fitted by restricted maximum likelihood, containing all 310 construct assays and all 73 wild-type assays together. Assay date entered as a random intercept, absorbing the day-to-day variation in the assay; a linear term in elapsed day absorbed the slower drift in wild-type fitness over the experiment. Fixed effects were the technician, an intercept for each recoded gene, a separate extent-of-recoding slope for each gene, and contrasts for recoding strategy with shuffling as the reference. Extent was expressed both as the fraction of a gene’s codons changed and as the number of codons changed per 100; both are reported.

Including the wild-type assays is what makes the technician term estimable. Wild type was assayed by both technicians throughout, including seven days on which both did so, and it carries no recoding treatment, so it identifies the technician difference independently of the strategy contrasts. In a model fitted to the recoded constructs alone the two are nearly aliased, because each technician assayed an almost disjoint set of strategies, and the separate effects are not identifiable. We also fitted a model allowing the technician effect to differ between wild type and recoded constructs, to test whether a technician difference could masquerade as a treatment effect.

Gene-specific slopes were tested by a likelihood ratio comparing a model with one slope per gene to a model with a common slope. Construct-to-construct variation was estimated by adding construct identity as a random effect nested within treatment (gene × strategy × level), so that its variance component measures variation among independently designed constructs sharing a treatment while the residual measures variation among repeated assays of the same construct. In gene F, the stable and unstable groups were compared by Welch’s t-test, and fitness was regressed on predicted folding energy within the ten mRNA folding constructs, which require no covariate because they change the same number of codons and share wild-type codon composition. The same regression was repeated across all 22 shuffled gene F constructs with the number of codons changed as a covariate, and across the twelve shuffled constructs that were not selected on folding energy. No comparison of folding energies was made between genes, because the achievable range differs by gene and the sequences were selected within genes. Differences in construction failure rates among genes were tested by Fisher’s exact test, standard logistic regression being inestimable because no gene F construct failed. The tables report descriptive means expressed relative to the same-day wild-type control; all inferential statements come from the model described above.

Overlap in edited codons between replicates was summarised as the number of codons that two replicates of the same treatment both edit, expressed as a percentage of the mean number each edits, and compared with the percentage expected if each had drawn the same number of codons at random from the pool editable by that strategy. The editable pool for each gene and strategy was taken as the number of distinct codons edited by at least one construct of that gene and strategy across all recoding levels; for the worst and random strategies this recovers the theoretical maxima given in the note to Table 1. The analysis is restricted to the worst, random and shuffle trios, which were selected for mutual divergence, and excludes the mRNA folding constructs, which were selected on a different criterion.

## Data availability

Raw sequencing reads are deposited in NCBI SRA (BioProject PRJNA1548792; run accessions SRR41316989-SRR41317006). Construct sequences, fitness measurements, and analysis code are available at https://github.com/jtvanleuven/phix174_recoding_code.

